# Identification of SMARCA1 as a key regulator for Colorectal Cancer

**DOI:** 10.1101/2024.12.24.630213

**Authors:** Chunqing Fu, Shoufeng Duan, Chengwen Zhang, Xinyu Cui, Fengxian Wang, Lichen Wang, Jinglei Hu, Tengjiao Li, Lin Li

## Abstract

Emerging evidence indicates that aberrations in the writing, reading, or erasure of chromatin modification codes are pivotal events in various human cancers. In this study, we conducted a systematic investigation of histone modification recognition proteins in a pair of colorectal cancer cell lines, SW480 and SW620. Using chromatin fractionation combined with data-independent acquisition mass spectrometry (DIA-MS), we developed a robust method to identify changes in histone modification recognition proteins during cancer progression. Our analysis revealed 22 proteins that were significantly upregulated and 22 proteins that were significantly downregulated in SW620 cells compared to SW480 cells. Notably, SMARCA1, a member of the ISWI family belonging to ATP-dependent chromatin remodeling complexes, was downregulated in SW620 cells compared to SW480 cells. Its high expression was strongly correlated with poor patient prognosis, aligning with the proposed role of SMARCA1 in promoting colorectal cancer (CRC) metastasis. Furthermore, reduced SMARCA1 expression altered the levels of metastasis-related matrix metalloproteinases (MMPs) in these cells. In conclusion, by systematically profiling histone modification recognition proteins in a matched pair of primary and metastatic CRC cell lines, we identified SMARCA1 as a potential driver of CRC metastasis and a promising therapeutic target for CRC patients.

## Introduction

Colorectal cancer (CRC) ranks as the third most prevalent cancer worldwide. While the incidence rate among older populations has declined in high-income economies, it is rising in emerging economies, particularly among individuals under the age of 50. A deeper understanding of the mechanisms underlying CRC metastasis could facilitate the development of more targeted treatments, ultimately improving patient outcomes.

Epigenetic alterations, including DNA methylation and demethylation, histone post-translational modifications and histone variants, ATP-dependent chromatin remodeling complexes, and non-coding RNA, play crucial roles in tumor initiation and progression [1]. Cancer cells exhibit complex and diverse behaviors driven by the coordinated expression of multiple genes, often diverging significantly from the gene expression patterns of their tissue of origin. This phenotypic plasticity underpins critical processes such as epithelial-to-mesenchymal transition (EMT), drug resistance, and enhanced proliferation. Recent research analyzing 1.7 million cells from 225 primary and metastatic tumor samples across 11 cancer types has identified DNA-level epigenetic changes, such as increased accessibility of regulatory subregions, as potential drivers of cancer progression [2]. These findings highlight the importance of deciphering the relationship between epigenetic states and gene expression regulation to better understand cancer phenotypic plasticity.

The SW480 and SW620 cell lines are derived from the primary tumor and its corresponding lymph-node metastatic tumor, respectively, from the same CRC patient, making them a valuable model for studying the genetic and molecular changes associated with CRC progression [3]. Xenograft assays have shown that SW620 tumors are more invasive and exhibit greater infiltration into adjacent normal tissues compared to SW480 tumors, although SW480 cells demonstrate higher migration and invasion capacities *in vitro* [3, 4]. Additionally, these two cell lines exhibit distinct morphologies in culture: SW480 cells display a spreading, epithelial-type morphology. These findings highlight significant phenotypic differences between SW480 and SW620, although the specific molecular mechanisms underlying these differences remain unclear. Top-down proteomic, targeted proteomics and isotope-labeled proteomics have been used to analyze protein expression differences including cellular secretion, Small GTPases and whole cell proteins between these two cell lines [5–8]. However, epigenetic differences between SW480 and SW620 cells have yet to be explored.

In this study, we performed a comparative analysis of histone modification recognition proteins in SW480 and SW620. Through bioinformatic analyses and experimental validation, we identified SMARCA1 as a key regulator of metastasis-related matrix metalloproteinases (MMPs) in CRC cells. Our findings suggest that SMARCA1 influences CRC cell migration and invasion and could serve as a promising therapeutic target for CRC patients.

## Results

### Quantitative profiling of differential expression of histone modification recognition proteins during CRC cell metastasis

To investigate critical factors involved in CRC cancer development, we established a proteomic workflow to quantitatively profile changes in histone modification recognition proteins in SW480 and SW620 cells. Our analysis focused on 218 histone modification recognition proteins. Using data-dependent acquisition (DDA)-based proteomics, we detected spectra corresponding to 79 proteins. To enhance proteome coverage, we constructed a coding sequence (CDS) library for the remaining 63 proteins. The final spectral library encompassed 177 proteins, each represented by one or two unique peptides.

Most studies quantify overall protein expression levels at the cellular level, which primarily reflect protein abundance in the cytoplasm or nucleoplasm. However, our approach focused specifically on functional proteins bound to chromatin. To this end, we performed chromatin fractionation on SW480 and SW620 cell extracts prior to proteomic analysis, enriching for chromatin-bound proteins (**Supplementary Figure S1A**).

Using the curated spectral library, data-independent acquisition mass spectrometry (DIA-MS) analysis enabled the quantification of 177 protein, covering more than half of the proteins in the library. In contrast, LC-MS/MS analysis in DDA mode identified only 79 of the 177 proteins in the chromatin-enriched fraction. This highlights the superior sensitivity of our DIA-MS-based approach compared to traditional shotgun proteomics.

### Differential Protein Expression in the CRC cell lines

Principal component analysis and correlation cluster analysis showed that DIA data of SW480 and SW620 were highly similar within their respective groups (**Figure 1A** and **1B**). Among the 177 quantified histone modification recognition proteins, 22 were significantly upregulated, and 22 were significantly downregulated (fold change >1.3, *p* < 0.05) in metastatic (SW620) cells compared to primary (SW480) CRC cells. These differences are visualized in the volcano plot and heatmap (Figure1C-1D). GSEA enrichment analysis (**Figure 1E**) showed that these genes were involved in ATP-dependent chromatin remodeling and transcriptional regulation of RNA polymerase II, etc. Their protein interaction network and functions were shown in **Figure 1F**, showing that they were mainly involved in histone methylation, demethylation, acetylation, deacetylation, phosphorylation, ubiquitination, deubiquitination, and chromatin remodeling as members of the SWI/SNF and ISWI families [9]. Notably, several histone modification recognition proteins, including upregulated proteins (HDAC8, KAT6A, MDC1, SMYD2, MRGBP, BRD7, CHEK1, BAZ1A, BPTF, HDAC1, ARID2 and KMT5A) and downregulated proteins (SMARCA2, KDM5B, SMARCA1, SETD7, ATR, BRCA1, EHMT1 and BARD1), exhibited pronounced expression changes (fold change > 1.5). In colorectal cancer case studies, SMARCA2, a member of the SWI/SNF complex, has been found to produce microsatellite instability in its absence [10]. HDAC8, a histone deacetylase, promotes CRC cell growth by downregulating IRF1, thereby modulating autophagy and facilitating liver metastasis [11]. KDM5B (Lysine-specific demethylase 5B) is overexpressed in various malignancies, including colorectal cancer (CRC). Elevated KDM5B expression has been observed in CRC tumor tissues compared to normal colon samples, with a positive correlation to cancer progression [12]. HDAC1, a histone deacetylase, has been reported to be upregulated in CRC tumors and is able to promote tumor angiogenesis through activation of the HIF1α/VEGFA signaling pathway [13]. SETD7, a SET domain-containing lysine methyltransferase, is significantly associated with tumor stage and microsatellite instability. Knockdown of SETD7 suppresses cancer cell proliferation in CRC [14]. Additionally, women with BRCA1 or BRCA2 mutations exhibit a fivefold increased risk of developing CRC [15].

**Figure 1.**
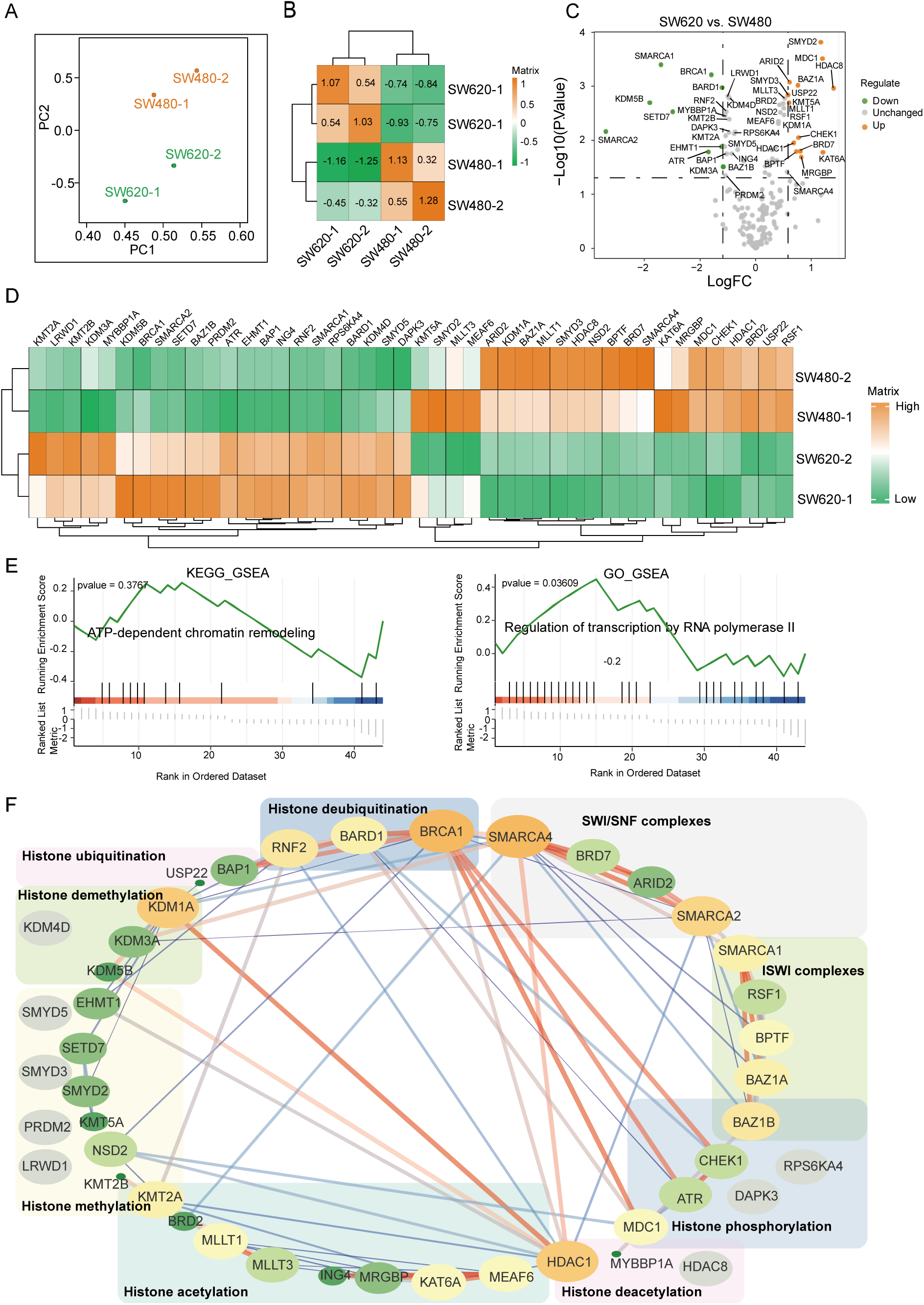
Quantitative profiling of differential expression of histone modification recognition proteins between SW480 and SW620 CRC cell lines. (A) Principal component analysis of histone modification-related proteins contained in colorectal cancer cell lines SW480 and SW620 by DIA-based mass spectrometry. **(B)** Pearson correlation heatmap of chromatin-associated proteins in SW480 and SW620 cell lines. **(C)** Chromatin-associated proteins with changes in SW620 relative to SW480 cell line using limma packages in R. The identification criteria were |logFC| > 0.3785, *p* value < 0.05. **(D)** Clustering analysis of differentially expressed chromatin-associated proteins between SW480 and SW620. The clustering algorithm was Euclidean distance. **(E)** GSEA enrichment analysis and signaling pathway analysis of differentially expressed proteins using the clusterProfiler package in R. The p value was set to 1. ATP-dependent chromatin remodeling was the only signaling pathway enriched. Regulation of transcription by RNA polymerase II was the biological process with the smallest pvalue value enriched. **(F)** Protein interaction analysis of differentially expressed proteins using the String database with a confidence level of 0.7. Larger shapes indicated higher degree values and more proteins associated with the protein. Thicker lines indicated higher combined scores.

SET and MYND domain-containing protein 2 (SMYD2) is highly expressed in multiple cancers, including CRC, where it activates the Wnt/β-catenin pathway and induces epithelial-mesenchymal transition (EMT) [16]. BRD7, a bromodomain-containing protein, acts as an oncogene in CRC, enhancing tumor progression by stabilizing c-MYC through ubiquitin–proteasome pathways [17]. Our findings corroborate these studies, reinforcing the oncogenic roles of SMARCA2, HDAC8, KDM5B, HDAC1, SETD7, SMYD2, BRD7, and other tumor-associated regulatory proteins in CRC progression.

### Potential Roles of SMARCA1 in CRC Progression

Among the differentially expressed proteins identified in our proteomic analysis, SMARCA1 (also known as SNF2L) emerged as a protein of interest, despite being relatively understudied. SMARCA1, a member of the ISWI family of chromatin remodeling proteins [18], is primarily known for its role in neurogenesis. In this study, we observed a significant downregulation of SMARCA1 in metastatic SW620 cells compared to primary SW480 cells (**Figure 1C-1D**, **Figure 2A, Supplementary Figure 1B**). This downregulation was further validated through western blotting, which corroborated the proteomic findings (**Figure 2B**).

**Figure 2.**
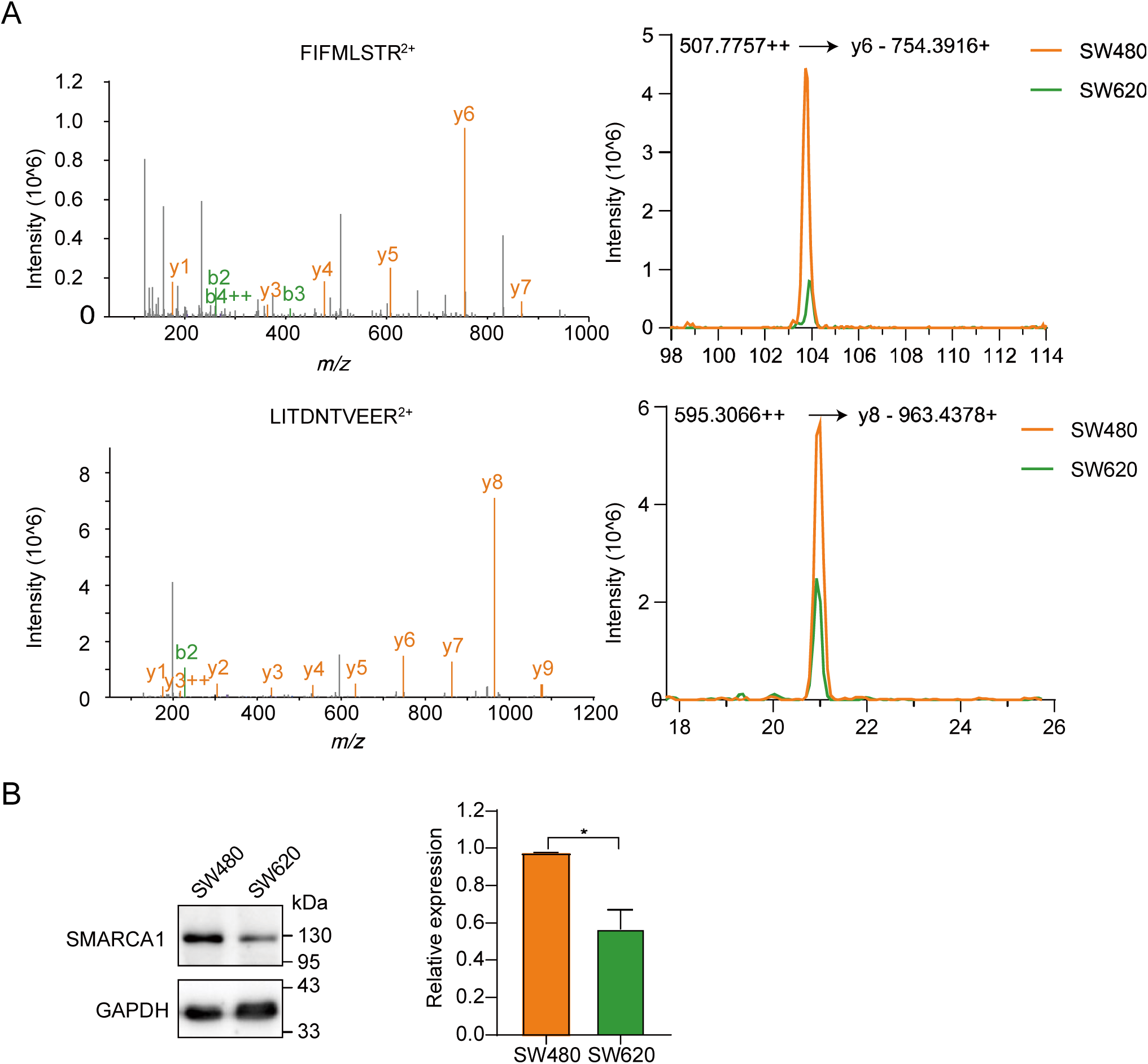
SMARCA1 was identified as a differentially expressed protein between SW480 and SW620 cell lines. **(A)** MSMS spectra of two peptides used to quantify SMARCA1 and their chromatograms in SW480 and SW620, respectively. **(B)** Differential expression of SMARCA1 in SW480 and SW620 detected using western blotting. *, *p* < 0.05.

To evaluate the prognostic significance of SMARCA1 in colorectal cancer (CRC), we analyzed patient cohorts from the TCGA-COAD and GSE21510 datasets (Figure 3A). Kaplan-Meier survival analysis revealed that elevated SMARCA1 expression was associated with a significantly increased risk of death [hazard ratio (HR) = 1.45, *p* = 0.00044, Log-rank test] and recurrence [hazard ratio (HR) = 1.43, *p* = 0.0013, Log-rank test] (**Figure 3A**). Additionally, SMARCA1 mRNA expression demonstrated a positive correlation with advancing CRC stages (**Figure 3B**). These findings underscore the potential role of SMARCA1 as a tumor driver and prognostic biomarker in CRC progression.

**Figure 3.**
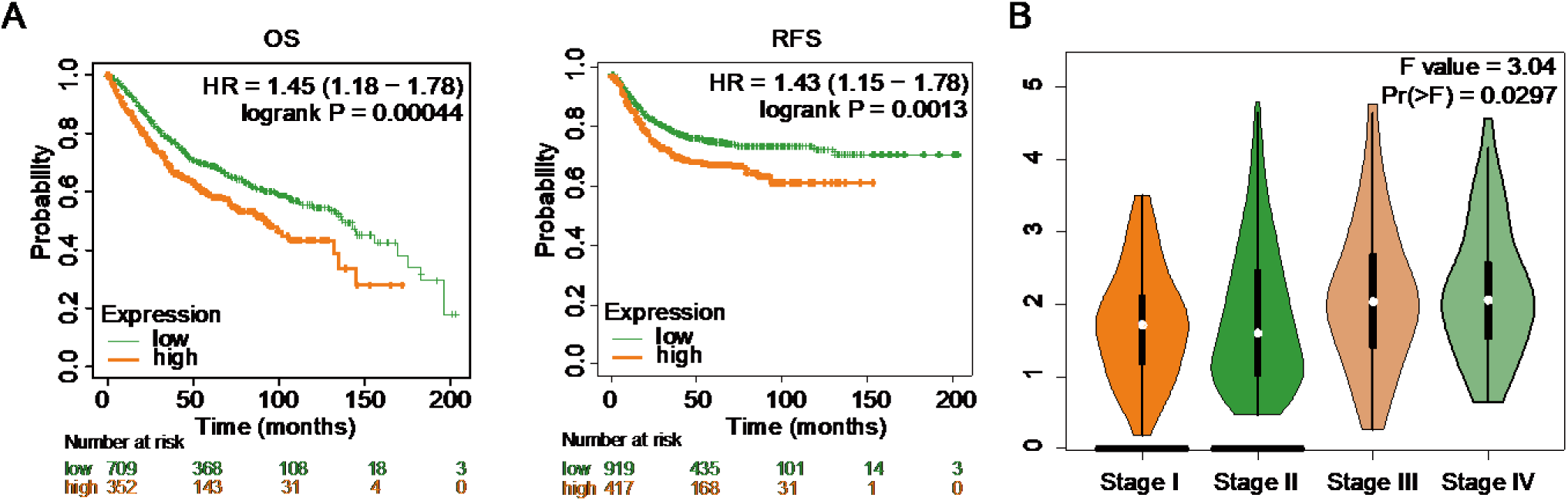
SMARCA1 influences the progression of colorectal cancer in patients. **(A)** Kaplan–Meier survival analysis of CRC patients stratified by the median SMARCA1 mRNA expression levels. OS, Overall survival. RFS, Recurrence free survival. **(B)** Differential mRNA expression of SMARCA1 in colorectal cancer patients across different stages of progression. Pr(>F), *p* value of the F statistic.

### Knockout of SMARCA1 Decreases *In Vitro* Migration and Invasion of SW480 Cells

SMARCA1 is broadly expressed in primary human tissues and has been reported to play either oncogenic or tumor-suppressive roles depending on the tumor type [18]. To investigate the functional role of SMARCA1 in CRC metastasis, we performed knockout experiments in SW480 cells. Silencing SMARCA1 did not significantly affect cell proliferation (**Figure 4A-4B**). However, it markedly enhanced the migratory and invasive abilities of SW480 cells, as evidenced by wound-healing and transwell invasion assays (**Figure 4C-4D**). These findings reinforce the involvement of SMARCA1 in regulating colon cancer progression and its potential role in CRC metastasis.

**Figure 4.**
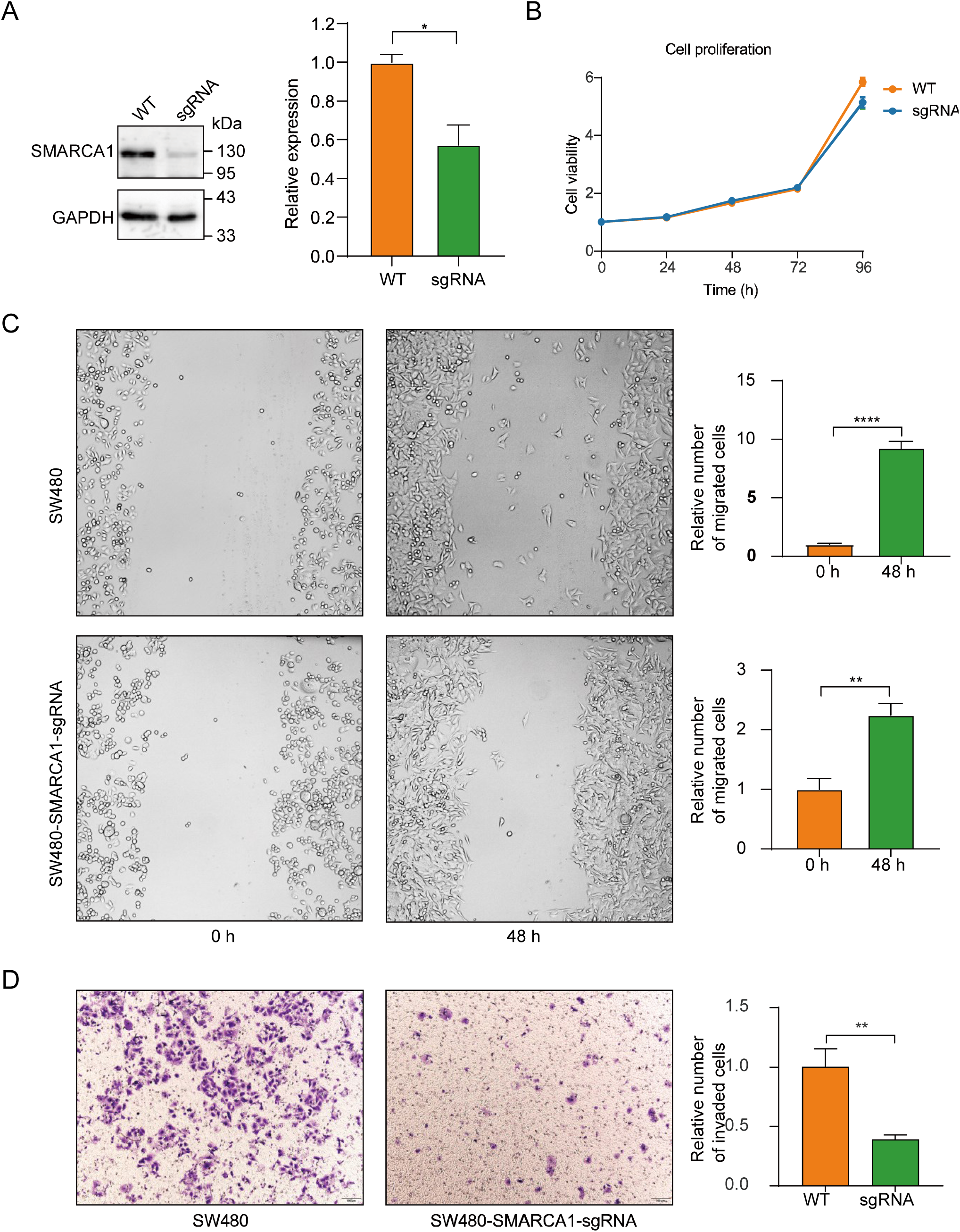
Knockout of SMARCA1 reduced in vivo migration and invasion in SW480 cells. **(A)** SMARCA1 knockout in SW480 cells was achieved using the CRISPR/Cas9-sgRNA system and verified by Western blot analysis. **(B)** Cell proliferation of wild-type and SMARCA1-knockout SW480 cells was measured using the CCK-8 assay. **(C)** Cell migration of wild-type and SMARCA1-knockout SW480 cells was assessed using the cell scratch assay. **, *p* value < 0.01. ****, *p* value < 0.001. **(D)** Cell invasion of wild-type and SMARCA1-knockout SW480 cells was evaluated using the transwell assay. **, *p* value < 0.01.

### Knockout of SMARCA1 Reduces the Expression of Key Metalloproteinases MMP2 and MMP14 While Increasing MMP7 in CRC Cells

The extracellular matrix (ECM), a critical component of the tumor microenvironment (TME), is extensively remodeled during cancer progression. Matrix metalloproteinases (MMPs) play pivotal roles in tumor invasion and metastasis by degrading the ECM [19]. Given the significant effects of SMARCA1 knockout on cell migration and invasion, we investigated its relationship with MMP expression. CRISPR/Cas9-mediated knockout of SMARCA1 in SW480 cells revealed decreased expression of MMP2 and MMP14, alongside elevated levels of MMP7 (**Figure 5A**). Consistently, when comparing SW480 and SW620 cells, the metastatic SW620 cells, which express lower levels of SMARCA1, exhibited reduced expression of MMP2 and MMP14 but increased expression of MMP7 (**Figure 5B**). These observations align with the effects seen in SMARCA1-knockout SW480 cells.

**Figure 5.**
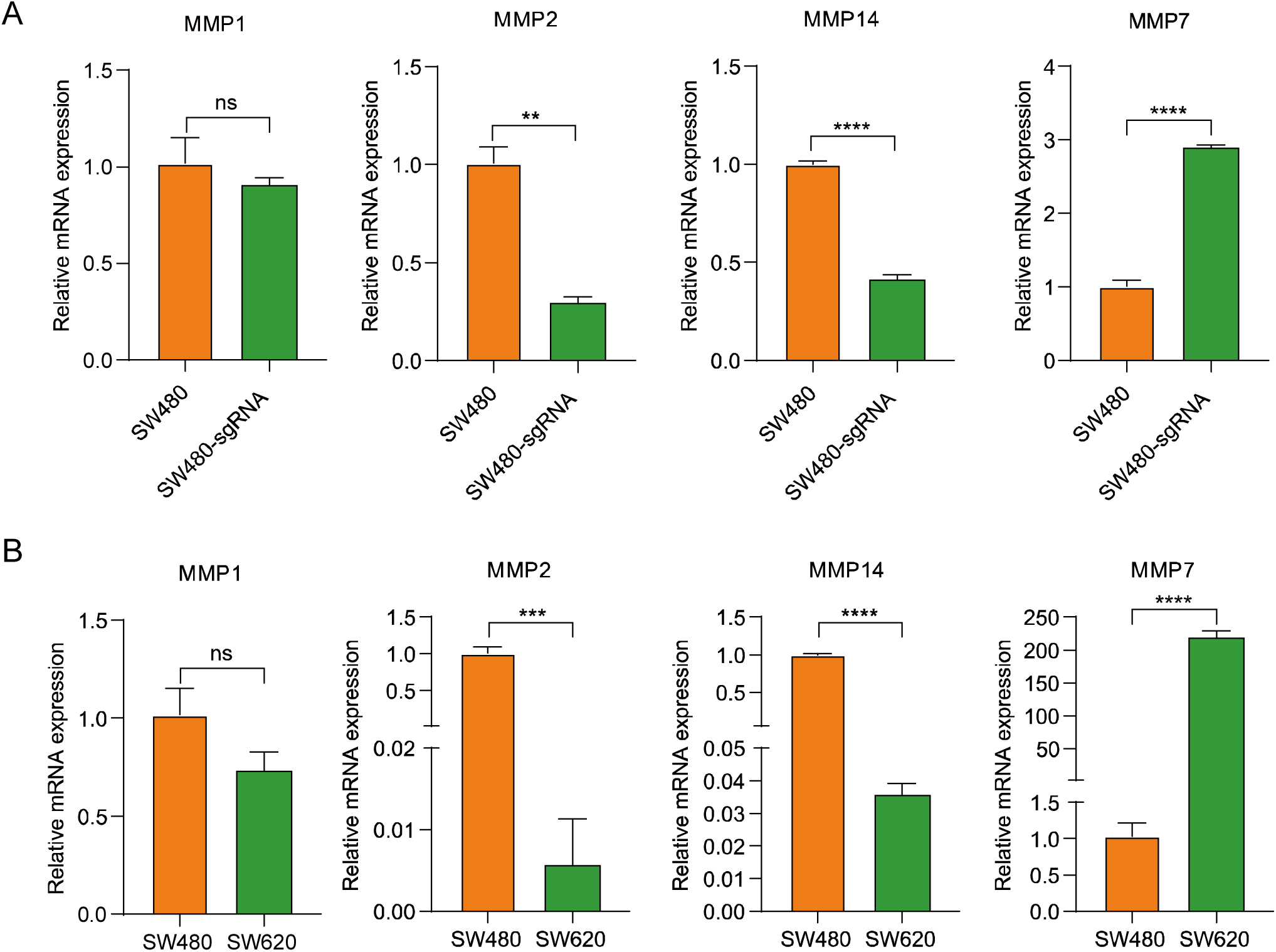
Knockout of SMARCA1 reduces the expression of key metalloproteinases MMP2 and MMP14, while increasing MMP7 in CRC Cell. **(A)** Differential mRNA expression of MMP1, MMP2, MMP14, and MMP7 in wild-type and SMARCA1-knockout SW480 cell lines. **(B)** Differential mRNA expression of MMP1, MMP2, MMP14, and MMP7 in SW480 and SW620 cell lines. ns, not significant; *, *p* value < 0.05; **, *p* value < 0.01; ***, *p* value < 0.005; ****, *p* value < 0.001.

Previous studies in various cancers have established a correlation between the proteolytic activity of MMP14 and activated MMP2 with endothelial cell invasion and pericellular ECM degradation. These processes are associated with angiogenesis and metastasis, underscoring their potential as therapeutic targets [20]. Conversely, MMP7 expression increases in later stages of colon cancer and correlates with poor prognosis [21, 22]. Our findings suggest that during CRC progression, SMARCA1 regulates the expression of distinct MMPs, which may contribute differentially to various stages of cancer development. These results highlight the role of SMARCA1 as a modulator of the tumor microenvironment and its potential as a therapeutic target in CRC.

## Discussion

In this study, we quantified 126 histone modification recognition proteins in the SW480/SW620 CRC cell line pair using chromatin fractionation combined with DIA-MS analysis. Unlike most previous studies that primarily focus on overall protein expression levels, we emphasized the importance of protein localization within the cell. Protein functionality is inherently tied to its proper cellular localization; proteins mislocalized or confined to incorrect compartments may lose their biological function or exert aberrant effects. Moreover, proteins at distinct locations can exhibit diverse biological roles. Ignoring these nuances could lead to misinterpretations in diagnostics or therapeutic target identification. By isolating chromatin-bound proteins, we aimed to enhance the precision and relevance of our epigenetic analyses.

Chromatin remodeling complexes contain a series of ATP-dependent remodeling enzymes that act as “molecular motors” to alter the interactions between DNA and histone octamers, thereby regulating the accessibility of DNA to transcription factors or other DNA-binding proteins [23]. The imitation switch (ISWI) family is a class of evolutionarily conserved ATP-dependent chromatin remodeling complexes that exhibit a wide range of aberrant gene expression and genetic status in human cancers, and alterations in components of the complexes are critical for tumorigenesis and progression [18].

SMARCA1, a member of the ISWI family, demonstrates context-dependent roles in cancer. It can act as either an oncogene or a tumor suppressor depending on the tumor type. For instance, SMARCA1 promotes cell survival and proliferation in many tumors [24]. Conversely, in gastric cancer, SMARCA1 knockdown inhibits tumor growth and disrupts cell homeostasis [25]. SMARCA1 expression is higher in normal melanoma cells than in malignant ones, and its depletion leads to increased cell proliferation and migration [26]. In CRC, it has been reported that long non-coding RNA DLEU1 in CRC cells can recruit SMARCA1 to the promoter of the KPNA3 and activate the expression of KPNA3, which promotes cell proliferation, migration and invasion of tumors [27]. And our study demonstrated that SMARCA1 can directly affect the migration and invasion of CRC cells, thus influencing tumor progression.

Our profiling of histone modification recognition proteins identified SMARCA1 as an oncogenic driver, enhancing cell migration and invasion (**Figure 4C-4D**). Kaplan–Meier survival analyses further substantiated its role, revealing a significant correlation between high SMARCA1 expression and poor patient outcomes (**Figure 3A**). Mechanistically, SMARCA1 knockout in SW480 cells resulted in decreased expression of MMP2 and MMP14, alongside increased MMP7 expression (**Figure 5**). This suggests that SMARCA1 influences the tumor microenvironment and metastatic potential of CRC cells through modulation of matrix metalloproteinases. However, the precise molecular pathways through which SMARCA1 contributes to CRC progression require further investigation.

In summary, we developed a robust proteomic workflow to profile histone modification recognition proteins, applying it to the SW480/SW620 CRC cell line pair. Among the identified proteins, SMARCA1 emerged as a key oncogene in CRC, promoting cell migration and invasion. Our findings suggest that SMARCA1 holds significant potential as a therapeutic target for CRC treatment, warranting further exploration into its mechanistic roles in tumor progression.

## Methods

### Cell Culture

SW480, SW620 and HEK293T cell lines were maintained in Dulbecco’s Modified Eagle Medium (DMEM, Thermo Fisher Scientific) supplemented with 10% fetal bovine serum (FBS, Adamas life, China) and penicillin/streptomycin (PS, 100 unit/mL) in a humidified atmosphere with 5% CO2 at 37 °C, and the culture medium was changed in every 2 to 3 days as necessary.

### Plasmid construction

In order to establish a mass spectrometric library of low-abundance proteins, the full-length complementary human cDNAs of ATAD2, BRD7, BRD1, BRPF1, PBRM1, DPF3, KAT6B, MLLT3, ING4, ATR, UBE2B, RNF2, BAP1, L3MBTL1, POLL, KDM3A, KDM4C, KDM4D, KDM5A, KDM5C, PSMA1, DOT1L, KMT5A, SETMAR, SMYD2, SETD7, PRDM3, CCNY, PAXIP1, SGF29, EHMT2, BDNF, CBX7, NBN, ORC1, JMJD6, PARP1, NEDD8, KAT8, HDAC3, SIRT1, USP3, USP15, USP22, BRCA1, BARD1, JAK2, RNF20, RNF40, ZMYND11, KDM3B, KDM5D, SETDB1, KDM6A, CARM1, MORF4L1, SIRT6, SIRT3, SIRT7, SIRT2, HDAC4, PHF8, and METTL9 were amplified using PCR technique and then cloned into the pFL3-flag vector, separately.

CRISPR-Cas9 system was carried out to decrease protein expression of SMARCA1. Three sgRNA sequences targeting SMARCA1 designed by CHOPCHOP (http://chopchop.cbu.uib.no/) and SYNTHEGO (https://design.synthego.com) website were 5’-GATAGTGGCGGTCGCATCCG-3’ (PAM CGG), 5’-GGATGCGACCGCCACTATCG-3’ (PAM TGG), and 5’-TTCAAATCTCTTTGCTCGGT-3’ (PAM CGG). These sequences were cloned into the pLV3-U6-MCS-sgRNA-Cas9-EGFP-Puro vector. Three plasmids were co-transfected into SW480 cell using Lipo8000 (Beyotime, China) in accordance with the protocol. Transfected cells were screened using 2 μg/mL puromycin to obtain the SW480 cell line with stable knockout of the SMARCA1.

### Expression and purification of low abundance histone modification-associated proteins

To obtain a mass library of low abundance proteins, each overexpression pFL3-flag plasmid containing CDS sequence was transfected into HEK293T cells using Lipo293 transfection reagent (Beyotime, China). All cells were collected after 48 h, and proteins were extracted using RIPA lysis buffer (50 mM Tris (pH 7.4), 150 mM HCl, 1% TrionX-100 (v/v), 1% sodium deoxycholate (w/v), 0.1%SDS (w/v), and protease inhibitor cocktail). Flag-tagged proteins were immunoprecipitated using anti-flag magnetic beads (SB-PR002, ShareBio, China) according to manufacturer’s instructions to obtain pure low-abundance proteins.

### Chromatin Isolation assay

To assess the differential expression of histone modification and chromatin remodeling related proteins in SW480 and SW620 cells, chromatin fraction was obtained with reference to the method of Juri Rappsilber’s group [28]. In brief, approximately 1 × 10^7^ cells were crosslinked using 1% formaldehyde, followed by termination of crosslinking using 0.25 M glycine. After PBS washing, cells were collected and lysed using cell lysis buffer (25 mM Tris (pH 7.4), 0.1% Triton X-100 (v/v), 85 mM KCl and protease inhibitor cocktail). Rnase A (Sangon, China) at 200 μg/ml was added to the solution to remove RNA. Cell nuclei were lysed using SDS buffer (50 mM Tris (pH 7.4), 10 mM EDTA, 4% SDS (w/v) and protease inhibitor cocktail) and solubility was increased using a 3-fold volume of urea buffer (10 mM Tris (pH 7.4), 1 mM EDTA and 8 M urea) to remove proteins not covalently bound to DNA. Subsequently, urea was washed using SDS buffer. Finally, the chromatin precipitate was resuspended using storage buffer (10 mM Tris (pH 7.4), 1 mM EDTA, 25 mM NaCl, 10% glycerol (v/v) and protease inhibitor cocktail), sonicated for 15 min at 10s-on,10s-off, 5% power (JY92-IIDN, ScientZ, China) to obtain chromatin proteins, and quantified using a BCA kit (ShareBio, China).

### Peptide Sample Preparation for Mass Spectrometry (MS)

For each sample, 100 μg of protein was mixed with SDS loading buffer (50 mM Tris-HCl (pH 6.8), 2% SDS (w/v), 10% Glycerol (v/v), 1% β-Mercaptoethanol (v/v), and 0.1% bromophenol blue (w/v)) and incubated at 95 °C for 30 min to reverse crosslink. All samples were run in 10% SDS-PAGE gels until the dye front was 1 cm from the bottom for higher reproducibility. The gels were washed with deionized water and decolorized using decolorizing solution (25 mM NH_4_HCO_3_ and 50% acetonitrile (ACN)). Each lane was cut into cubes of ≈ 1 mm^2^ and dehydrated with ACN for additional 10 min with shaking. Dehydrated gel pieces were reduced with 10 mM dithiothreitol (DTT, Adamas, China) at 37°C for 60 min and then dehydrated with ACN. Alkylation was conducted by replacing the DTT solution with 25 mM iodoacetamide (IAA, Sigma-Aldrich) and incubated at room temperature for 20 min in the dark followed by dehydrating with ACN. The gel pieces were digested with 2 ng/μL of trypsin (Thermo Fisher Scientific) in 25 mM NH_4_HCO_3_ overnight at 37 °C. After digestion, the supernatant was collected to obtain the peptides. To increase the yield of peptides, the supernatant containing peptides was collected by adding using 65% ACN and 5% formic acid (FA) to the gel and sonicated for 5 min in a water bath and incubated at 37°C for 30 min. ACN was added for the same extraction process as before and the supernatant was collected. All the supernatants were combined and dried in a vacuum centrifugal dryer (Thermo Fisher Scientific) at 50°C and resuspended in 100 μL of 0.1% formic acid. All samples were desalinated using Empore StageTips (6091, CDS Analytical, US).

### LC-MS/MS Analyses

The dried peptide samples were re-dissolved with 0.1% FA and further analyzed by electrospray liquid chromatography-tandem mass spectrometry using an Orbitrap Exploris 480 (Thermo Fisher Scientific) coupled to an EASY-nLC 1200 (Thermo Fisher Scientific). Samples were run on a C_18_ Column (3 μm particle size, 75 μm × 20 cm, 100 Å) at a flow rate of 250 nL/min using the following gradient: 0 to 5 min 2 to 8% B, 5 to 97 min 8 to 22% B, 97 to 110 min 22 to 35% B, 110 to 111 min 35 to 90% B, 111 to 120 min at 80% B. Mobile phase A was 0.1% FA in water and Mobile phase B was 0.1% FA in 80%ACN. DDA MS parameters were set to: (1) MS: scan range (m/z), 350-1,500; orbitrap resolution, 60,000; time (ms), 20; normalized AGC target (%), 300; number of dependent scans, 30; exclusion duration, 30 s; exclude after n times, 1. (2) HCD-MS/MS: orbitrap resolution, 15,000; time (ms), 20; normalized AGC target (%), 100; HCD CE (%), 30; include charge exclusion, 2-6; 1; 1.0 m/z isolation window. The same nano-LC system and gradient are used for DIA analysis. DIA MS parameters were set to: (1) MS: scan range (m/z), 350-1,500; orbitrap resolution, 60,000; ACG target, standard; cycle time (s), 3. (2) HCD-MS/MS: orbitrap resolution, 30,000; scan range (m/z), 200-2000, time (ms), 54; normalized AGC target (%), 1000; HCD CE (%), 32; 60 DIA isolation windows of 10 *m/z* from m/z 400-1000. The DDA data were searched using MaxQuant (version 2.4.9), and then the search data were imported into Skyline software (version 23.1) to create a mass spectral library of 271 histone modification and chromatin remodeling related proteins. The DIA data were imported into Skyline, and the chromatograms of the peptides of the related proteins were extracted for quantification and comparison.

To assess the overall variance of the data obtained from skyline analysis, principal component analysis was performed using fast.prcomp function in R. Correlation clustering of samples was performed using Pearson correlation coefficient. Differentially expressed proteins (DEP) were identified by limma package in R (version 4.3.0) with a cutoff of fold change >1.3 and *p* value <0.05, which is Characterized by volcano map. Cluster analysis of DEP was plotted using the pheatmap function in R and the clustering method was the Euclidean distance method. GSEA enrichment (GO) and signaling pathway (KEGG) analysis of DEP were performed using the clusterProfiler package in R. Due to the small number of DEP was very limited, *p* value was set to 1 to obtain more information. Protein interaction network analysis (PPI) of DEP was carried out using the STRING database with a confidence level set to 0.7 and the result was furthered visualized by Cytoscape (Version 3.10.3).

### Quantitative real-time PCR (RT-qPCR)

Eastep Super Total RNA Extraction Kit (LS1040, Promega) was employed to extract total RNA following manufacturer’s instructions. The mRNA levels was assessed using HiScript III RT SuperMix for qPCR (+gDNA wiper) (R323, Vazyme, China) and 2×Universal SYBR Green qPCR Premix (Q312-02, Vazyme, China). All results were normalized to GAPDH. The relative expression of mRNAs was quantified using the 2^−ΔΔCt^ method. Primer sequences used were: GAPDH, forward (F) 5’-GTCGGAGTCAACGGATTTGG-3’, reverse (R) 5’ -TGCCATGGGTGGAATCATATTG-3’; MMP1, F 5’-AGAGCAGATGTGGACCATGC-3’, R 5’-TTGTCCCGATGATCTCCCCT-3’; MMP2, F 5’-GTCCCCATGAAGCCCTGTTC-3’, R 5’-CCCTGGAAGCGGAATGGAAA-3’; MMP7, second round F 5’-AACAATTGTCTCTGGACGGC-3’, R 5’-TCTCTTGAGATAGTCCTGAGCCT-3’; MMP14, F 5’-GCGTCCATCAACACTGCCTA-3’, R 5’-CACCCAATGCTTGTCTCCTTTG -3’.

### Protein extraction and western blotting

Cells were directly harvested with SDS loading buffer (50 mM Tris-HCl (pH 6.8), 2% SDS (w/v), 10% Glycerol (v/v), 1% β-Mercaptoethanol (v/v), and 0.1% bromophenol blue (w/v)). Protein expression was detected by Western blotting. Equal amounts of proteins were fractionated by 12% sodium dodecyl sulfate (SDS)-polyacrylamide gel electrophoresis. Anti-SMARCA1 (A10248, ABclonal, China) and Anti-GAPDH (SB-AB0037, ShareBio, China) were used and the expression signal was amplified by incubating the secondary antibody of the corresponding species. The blots were detected using ECL chemiluminescence solution (SB-WB012, ShareBio, China) on a ChemiDoc Imaging Systems (BioRad). Grayscale values of the target bands were calculated using ImageJ for quantitative comparison.

### Wound healing assay

Differences in migration rates between wild type and SMARCA1-knockout SW480 cells were assessed using a wound healing assay. Cells were seeded and gently scraped with a 10 μL sterile pipette tip when the cells achieved 90% confluence. The wounded cells were continuously cultured for 48 hours. The healing situation were recorded at the 0 h and 48 h under the IX73 inverted fluorescence microscope (Olympus Corporation, Japan). The migration rate was measured by the ratio of the cell area of day 0 and day 2 measured by ImageJ.

### Transwell assays

To assess cell invasion, the transwell membrane (6.5 mm in diameter with 8.0 µm pores, Corning, USA) were coated with a 300 ng/µL Matrigel solution (ABW, 082704, Shanghai Nova Pharmaceutical Technology, China) according to manufacturer’s instructions. 1 × 10^5^ wild type and SMARCA1-knockout SW480 cells were seeded into the inner chamber in serum-free medium, and the outer chamber was filled with medium containing 10% FBS. After 72 h of incubation, non-invaded cells at the inner bottom of the chamber were scraped off using a cotton swab, and invaded cells at the outer bottom of the chamber were fixed with methanol for 15 min and stained with 0.1% crystal violet for another 15 min. Then 5 fields were selected and photographed using an BX53 upright fluorescence microscope (Olympus Corporation, Japan). The invasion differences of different cells at 72 h were evaluated by measuring the stained cell area using ImageJ.

### Online Data Analysis

Survival effects of SMARCA1 expression in CRC patients were analyzed using the Kaplan-Meier Plotter online website, including overall survival analysis (OS) and Recurrence free survival (RFS) [29]. Differential expression of SMARCA1 in CRC patients with different developmental stages was analyzed using GEPIA2 online platform (http://gepia2.cancer-pku.cn/#index) integrated with The Cancer Genome Atlas (TCGA) projects [30].

## Acknowledgments

This work was financially supported by the National Natural Science Foundation of China (Grant No. 22104084) and Startup funding from Shanghai Jiao Tong University (SJTU).

## Notes

### Competing Interest Statement

The authors have declared no competing interest.

