## Supplementary Figure S1 for "Identification of SMARCA1 as a key regulator for Colorectal Cancer"

**Supplementary Figure 1**

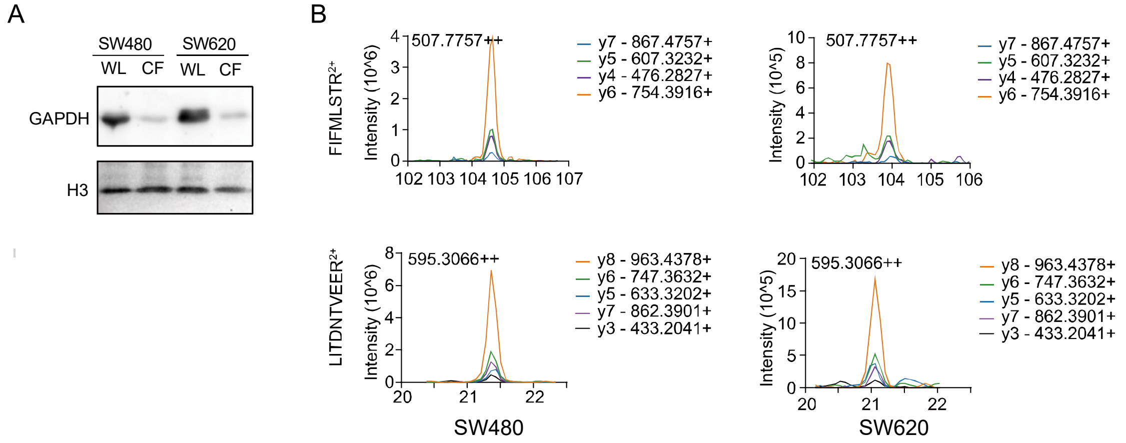


**Supplementary figure 1** Verification of chromatin fractionation in both SW480 and SW620 cell lines, and the chromatograms of SMARCA1 peptides.

**(A)** Chromatin fractionation efficiency was verified by Western blotting in SW480 and SW620. WL, whole lysate. CF, chromatin fraction. (B) Chromatograms of two peptides identified by DIA mass spectrometry in SW480 and SW620 cell lines, respectively.
